# LncRNAs Landscape in the patients of primary gout by Microarray Analysis

**DOI:** 10.1101/2020.04.27.063768

**Authors:** Yu-Feng Qing, Jian-Xiong Zheng, Yi-Ping Tang, Fei Dai, Zeng-Rong Dong, Quan-Bo Zhang

## Abstract

To determine the differential profiles of long non-coding RNAs (lncRNAs) in Peripheral blood mononuclear cells (PBMCs) among acute gout (AG) patients, intercritical gout (IG) patients and healthy control subjects, and to explore the specific biomarkers for acute gout diagnosis and treatment in future. Human lncRNA microarrays were used to identify the differentially expressed lncRNAs and mRNAs in primary AG (n=3), IG (n=3) and healthy subjects (n=3). For comparison Bioinformatics analyses were performed to predict the roles of abberrantly expressed lncRNAs and mRNAs. Quantitative real-time polymerase chain reaction (qRT-PCR) was applied to validate the results in 32 AG, 32 IG patients and 32 healthy control subjects. A receiver operating characteristic (ROC) curve was constructed to evaluate the diagnostic value of the lncRNAs identified in gout. The microarray analysis identified 3421 and 1739 differentially expressed lncRNAs, 2240 and 1027 differentially expressed mRNAs in AG and IG (fold change>1.5, P<0.05; respectively), respectively. qRT-PCR results of 9 dysregulated genes were consistent with the microarray data. The bioinformatic analysis indicated that the differentially expressed lncRNAs regulated the abnormally expressed mRNAs, which were involved in the pathogenesis of gout through several different pathways. The expression levels of TCONS_00004393 and ENST00000566457 were significantly increased in the AG group than those in the IG group or healthy subjects (P<0.01, respectively). Moreover, the areas under the ROC curve were 0.955 and 0.961 for TCONS_00004393 and ENST00000566457,respectively. Our results provide novel insight into the mechanisms of the primary gout, and reveal that TCONS_00004393 and ENST00000566457 might be as candidate diagnostic biomarkers and targets for the treatment of acute gout.

## Introduction

Gout, the one of most common form of autoinflammatory arthritis in human, is characterized by elevated urate and monosodium urate (MSU) crystal deposition in tissues,which leads to arthritis, occurrence of soft tissue masses (i.e., tophi), nephrolithiasis, and urate nephropathy [1]. The epidemiological evidence suggests that both the incidence and prevalence of gout are rising, and the incidence is 1.14% and 1.4% in Shandong coastal cities of Eastern China and eastern counties respectively [2].The specific pathogenesis of gout is still unclear. Previous studies have demonstrated that an attack of gouty arthritis is triggered by the deposition of MSU crystals in the joint and MSU crystals is widely recognized as endogenous danger signal by components of the innate immune system [3,4]. Hyperuricemia is the biochemical basis of gout [5].

Long noncoding RNAs (lncRNAs) are typified by a length of transcription longer than 200 nucleotides that is not translated into proteins [6].Increasing scientific interest in these factors stems from previous investigations showing that lncRNAs exert regulatory effects on gene expression levels, involving epigenetic regulation, transcriptional regulation, and post-transcritional regulation in the form of RNA[7]. LncRNAs also play important roles in modulating innate and adaptive immune responses and immune cell development [8]. Moreover, emerging evidence suggests that lncRNA could be as diagnostic biomarkers in many diseases such as some cancer[9], osteoporosis[10], Alzheimer’s disease[11], ect. So, lncRNAs have attracted much attention in recent medical studies.

The diagnosis of gout depends heavily on the higher concentration of uric acid in the blood. However, the concentration of blood uric acid were not observed increased in some acute gout patients[12]. So, the other serum biomarkers of diagnosis in acute gout are urgently needed. Few research of lcnRNA in gout are reported. Therefore, the present study was initiated to use lncRNA microarray for the characterization of genome-wide lncRNA and messenger RNA (mRNA) expression profiles of acute gout patients compared with intercritical gout patients and healthy control subjects. Our goal was to establish the potential utility of lncRNAs as biomarkers of diagnosis or treatment targets for acute gout.

## Materials and Methods

### Patients and sample collection

Sixty-four consecutive male patients with primary gout from the Department of Rheumatology and Immunology of Affiliated Hospital of North Sichuan Medical College between February 2019 and January 2020, were enrolled into this study. The classification of gout fulfilled the 1977 American Rheumatism Association (now the American College of Rheumatology (ACR)) preliminary criteria for the classification of the acute arthritis of primary gout, as well as the 2015 ACR/European League Against Rheumatism (EULAR) gout classification criteria[13,14]. All the gout patients had no history of cancer, hematopathy, nephropathy, infection or other autoimmune diseases. Gout patients were divided into an acute gout (AG) group (including 32 patients with acute gout attacks) and an intercritical gout (IG) group (including 32 patients) based on whether patients are presenting onset of symptoms or not.The gout patients were not receiving any systemic anti-inflammatory treatment, or drugs to control the production and elimination of uric acid before blood samples obtained. Healthy controls (32 age-matched men) with no hyperuricemia, no metabolic syndrome, and no other chronic diseases were recruited for the comparative study. The demographic and clinical features of all the subjects are summarized in Table 1. Peripheral blood anticoagulated with ethelene diamine tetraccetic (EDTA) was obtained from all subjects. Human peripheral blood mononuclear cells (PBMCs) were isolated with Ficoll-Hypaque gradients. Written informed consent was obtained from all of the enrolled participants.This study was approved by the Ethics Committee of the Affiliated Hospital, North Sichuan Medical College and conducted in accordance with the ethical guidelines of the 1975 Declaration of Helsinki.

**Table 1.**
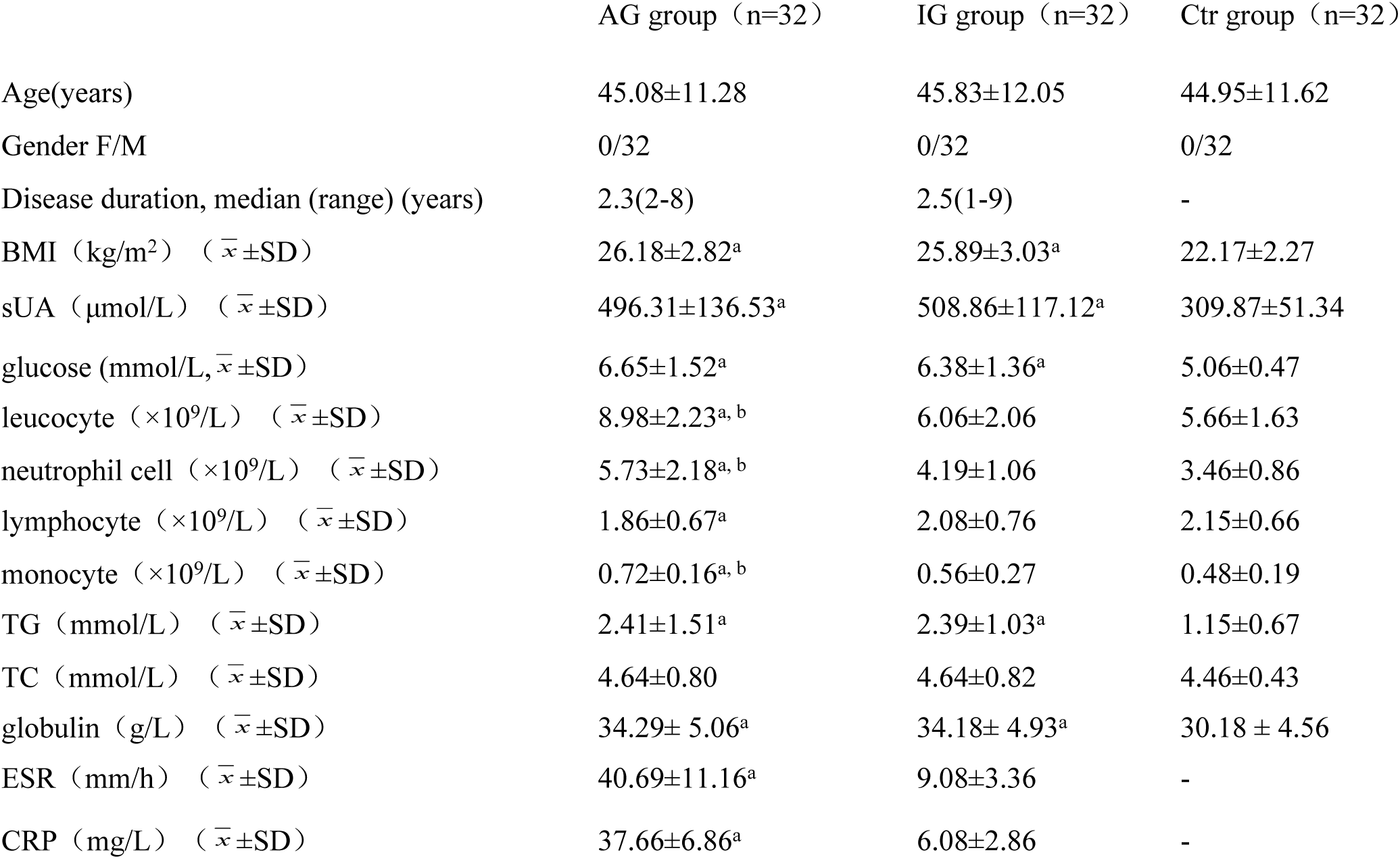
Clinical and laboratory data of subjects studied

|  | AG group (n=32) | IG group (n=32) | Ctrl group (n=32) |
| --- | --- | --- | --- |
| Age(years) | 45.08±11.28 | 45.83±12.05 | 44.95±11.62 |
| Gender F/M | 0/32 | 0/32 | 0/32 |
| Disease duration, median (range) (years) | 2.3(2-8) | 2.5(1-9) | - |
| BMI (kg/m<sup>2</sup>) ($\bar{x} \pm SD$) | 26.18±2.82<sup>a</sup> | 25.89±3.03<sup>a</sup> | 22.17±2.27 |
| sUA (μmol/L) ($\bar{x} \pm SD$) | 496.31±136.53<sup>a</sup> | 508.86±117.12<sup>a</sup> | 309.87±51.34 |
| glucose (mmol/L, $\bar{x} \pm SD$) | 6.65±1.52<sup>a</sup> | 6.38±1.36<sup>a</sup> | 5.06±0.47 |
| leucocyte (×10<sup>9</sup>/L) ($\bar{x} \pm SD$) | 8.98±2.23<sup>a, b</sup> | 6.06±2.06 | 5.66±1.63 |
| neutrophil cell (×10<sup>9</sup>/L) ($\bar{x} \pm SD$) | 5.73±2.18<sup>a, b</sup> | 4.19±1.06 | 3.46±0.86 |
| lymphocyte (×10<sup>9</sup>/L) ($\bar{x} \pm SD$) | 1.86±0.67<sup>a</sup> | 2.08±0.76 | 2.15±0.66 |
| monocyte (×10<sup>9</sup>/L) ($\bar{x} \pm SD$) | 0.72±0.16<sup>a, b</sup> | 0.56±0.27 | 0.48±0.19 |
| TG (mmol/L) ($\bar{x} \pm SD$) | 2.41±1.51<sup>a</sup> | 2.39±1.03<sup>a</sup> | 1.15±0.67 |
| TC (mmol/L) ($\bar{x} \pm SD$) | 4.64±0.80 | 4.64±0.82 | 4.46±0.43 |
| globulin (g/L) ($\bar{x} \pm SD$) | 34.29± 5.06<sup>a</sup> | 34.18± 4.93<sup>a</sup> | 30.18 ± 4.56 |
| ESR (mm/h) ($\bar{x} \pm SD$) | 40.69±11.16<sup>a</sup> | 9.08±3.36 | - |
| CRP (mg/L) ($\bar{x} \pm SD$) | 37.66±6.86<sup>a</sup> | 6.08±2.86 | - |

### RNA Extraction

>For RNA purification, we used TRIzol reagent (Invitrogen, Gran Island, NY, USA) according to the manufacturer’s instructions followed by application of PBMCs to RNeasy spin columns (Qiagen, Venlo, Limburg, Netherlands). RNA quantity and quality were measured by NanoDrop ND-1000. RNA integrity was assessed by standard denaturing agarose gel electrophoresis or Agilent 2100 Bioanalyzer.

### lncRNA Microarray

The Human LncRNA Microarray V4.0 (Arraystar,Rockvill,MD,USA) was used to design the global profiling of human lncRNAs and protein-coding transcripts, which is updated from the previous Microarray V3.0. The fourth-generation lncRNA microarray detects about 40,173 LncRNAs and 20,730 coding transcripts. The lncRNAs were carefully constructed using the most highly respected public transcriptome databases (RefSeq, UCSC Known Genes, and Genecode, ect.), as well as landmark publications. A specific exon or spice junction probe was used to represent each transcript that could accurately identify individual transcripts.

### qRT-PCR Validation

To validate the microarray data, we selected five lncRNs (NR_026756, ENST00000566457, TCONS_00004393, TCONS_00003286, and NR_029386) and four mRNAs (NFkBIZ, PPP4R3B, EGR3 and IL-1β) from aberrantly expressed lncRNAs and mRNAs.

Quantitative real-time reverse transcription PCR (qRT-PCR) is the gold standard for data verification in this context. For the reverse transcriptase (RT) reaction,SYBR Green RT reagents (Bio-Rad, USA) were used. In brief, the RT reaction was performed for 60 min at 37 ℃, followed by 60 min at 42 ℃, using oligo (dT) and random hexamers. PCR amplifications were performed by using SYBR Green Universal Master Mix. These reactions were performed in duplicate containing 2×concentrated Universal Master Mix, 1µl of template cDNA, and 100nM of primers in a final volume 12.5µl, followed by analysis in a 96-well optical reaction plate (Bio-rad). The PCR results were quantified by using the method 2^-ΔΔct^ method against β-actin for normalization. These data represent the means of three experiments.

### GO and Pathway Analysis

Gene ontology (GO) analysis (www.geneontology.org) was used to investigate biological functions based on differentially expressed coding genes[15]. This analysis classifies functions according to the three following aspects: biological process, cellular component and molecular function. Fisher’s exact test was applied to classify the GO category. The p-value denotes the significance of GO term enrichment in the deregulated expressed genes. The lower the p-value, the more significant the GO term (P value<0.05 is recommended).

Pathway analysis was used to investigate the differentially expressed coding genes according to the Kyoto Encyclopedia of Genes and Genomes (KEGG), Biocarta and Reactome (http://www.genome.jp/kegg/)[16]. The p-value (EASE-score, Fisher-P value or Hypergeometric-P value) indicates the significance of the pathway correlated with the conditions. P<0.05 was considered statistically significant.

### LncRNA-mRNA Co-Expression Network

The lncRNA-mRNA co-expression network identifies interactions between differentially expressed mRNAs and differentially expressed lncRNA. The basis of this construct is the normalized signal intensities of specific expression levels of mRNAs and lncRNAs. To formulate the lncRNA-mRNA co-expression network used here, we applied Pearson’s correlations, to calculate statistically significant associations.

### Statistical analysis

A one-way ANOVA test or unpaired t-test was used for statistical analysis. Receiver operating characteristic (ROC) analysis was used to evaluate the power of each candidate biomarker. All statistical tests were performed using SPSS 17.0. P-values of <0.05 were considered statistically significant.

## Results

### LncRNA and mRNA profiles differ among acute gout patients, intercritical gout patients and healthy control subjects (Table 2)

Human LncRNA Microarray V4.0 was used to detect lncRNAs in PBMCs from six gout patients (including three acute gout and three intercritical gout patients) and three healthy control subjects. Volcano plot analysis was then applied for the identification of differences in lncRNAs and mRNAs from three group populations (Fig 1). Cluster assays of the 1000 randomly selected lncRNAs and mRNAs indicated that the expression pattern of the acute gout (AG) group was quite different than those of the intercritical gout (IG) and healthy control groups (Fig 1). Compared with the healthy control group, the AG and the IG groups exhibited 2147 and 1046 up-regulated lncRNAs, respectively, whereas 1274 and 693 lncRNAs were expressed at a lower level in the AG and IG patients compared with the healthy control subjects (Table 2). In the AG group, compared with the IG group, 2855 lncRNAs were differentially expressed, including 1931 up-regulated and 924 down-regulated lncRNAs (Table 2). As shown in table1, total of 1240 mRNAS (836 up-regulated and 1404 down-regulated) were significantly aberrantly expressed in AG patients compared with those in healthy control subjects. The Venn diagram further indicated that the AG and IG groups shared 384 up-regulated and 423 down-regulated lncRNAs (Fig 2A), 196 up-regulated and 247 down-regulated mRNAs (Fig 2B)

**Table 2.** Number of differentially expressed lncRNAs and mRNAs in the different groups lncRNAS

|  | lncRNAs |  |  | mRNAs |  |  |
| --- | --- | --- | --- | --- | --- | --- |
|  | AG vs Ctrl | IG vs Ctrl | AG vs IG | AG vs Ctrl | IG vs Ctrl | AG vs IG |
| up-regulated | 2147 | 1046 | 1931 | 836 | 520 | 870 |
| down-regulated | 1274 | 693 | 924 | 1404 | 507 | 784 |

AG:acute gout, IG: intercritical gout, Ctrl:healthy control subjects

**Fig 1.**
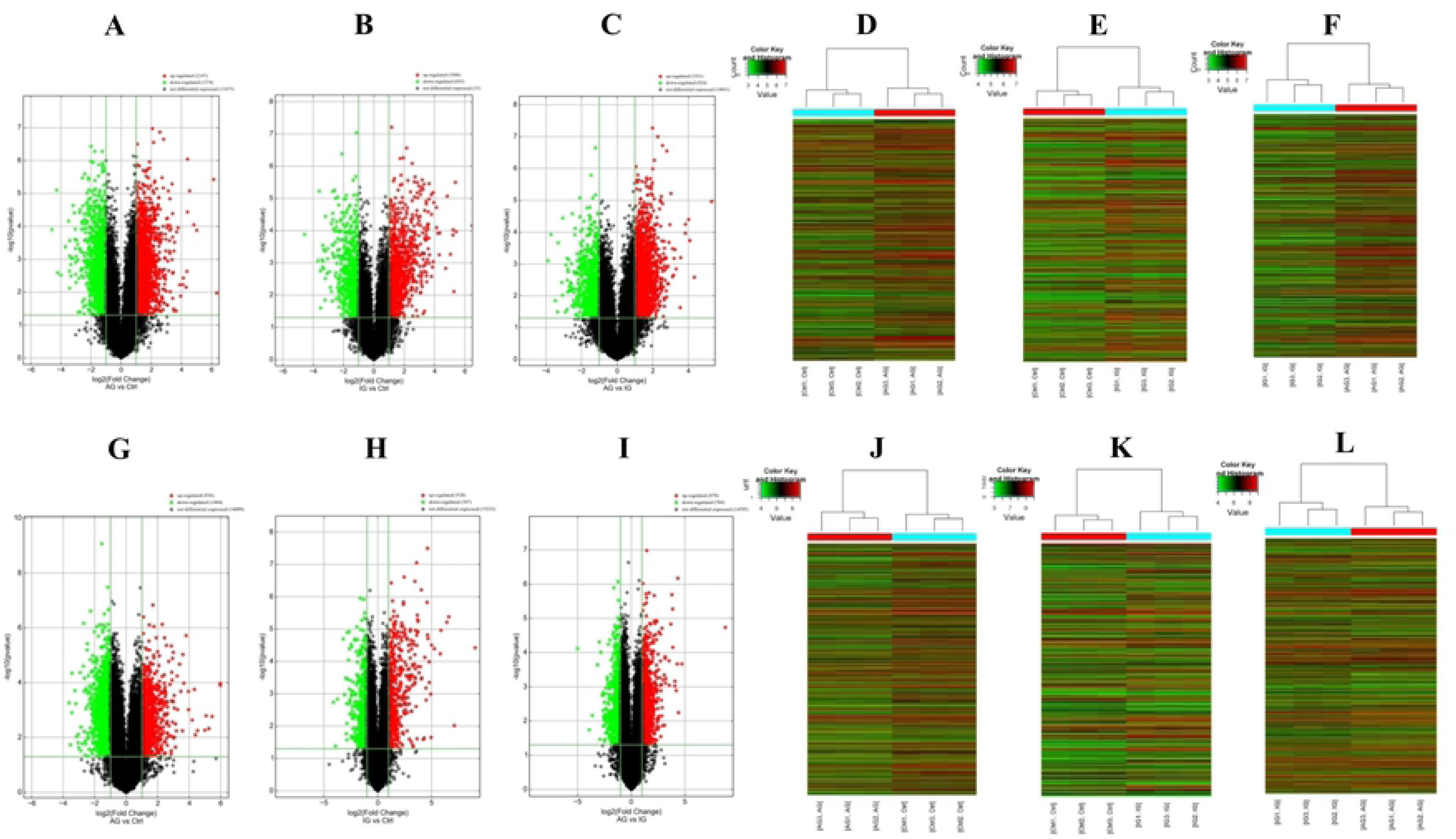
Expression profiles of lncRNAs(A-F) and mRNAs (G-L) in gout patients. **(A-C,G-I)** Volcano plot of differentially expressed lncRNAs and mRNAs. The horizontal green line represents a P-value of 0.05, and vertical green lines represent 2.0-fold changes up and down. X axes are the fold change values (log2 scaled), and Y axes are the P-values (log10 scaled). **(D-F,J-L)** Hierarchical clustering analysis of lncRNAs and mRNAs with expression changes greater than two-fold and P value <0.05. AG (acute gout) patients group: AG1, AG2, AG3; IG (intercritical gout) patients group: IG1, IG2, IG3; Ctr (healthy subjects) group: Ctr1, Ctr2, Ctr3. Red and green colors represent up- and down-regulated genes, respectively.

**Fig 2.**
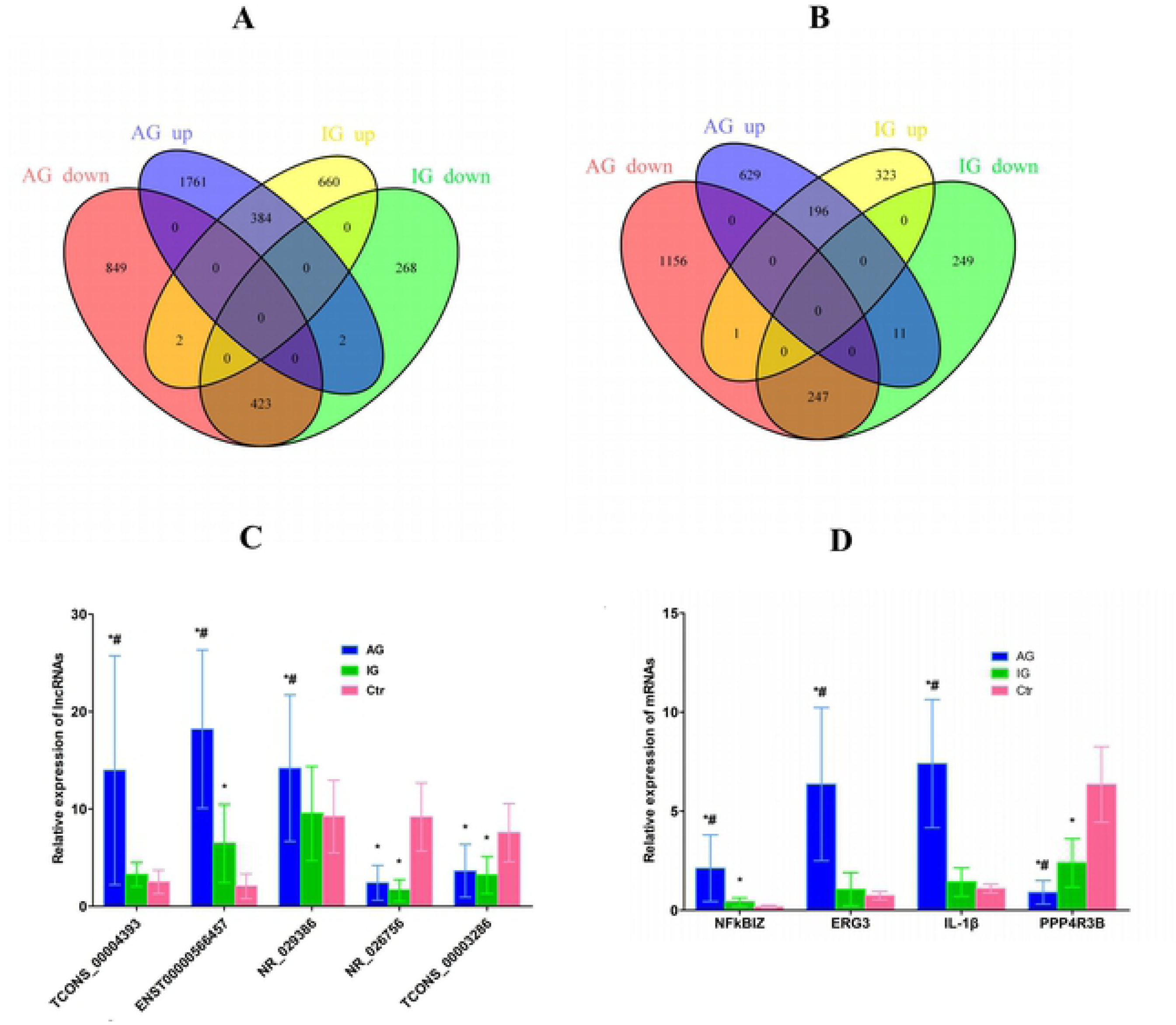
Differential expression of lncRNAs and mRNAs in PBMCs between the AG group, IG group and the healthy Ctr group. **A and B.** Venn diagrams indicated the numbers of overlapping and non-overlapping differentially expressed lncRNAs and mRNAs in the AG or the IG compared with the Ctr, respectively. **C and D.** Differential expression results of lncRNAs and mRNAs validated with qRT-PCR. * represents statistical significance (P<0.05) compared with the healthy control group, and # represents statistical significance (P<0.05) compared with the IG group. AG represents acute gout, IG represents intercritical gout.

### qRT-PCR Validation

To validate our results independently and determine the role of lncRNAs in gout patients, five aberrantly expressed lncRNAs (NR_026756, ENST00000566457, TCONS_00004393, TCONS_00003286, and NR_029386) and four aberrantly expressed mRNAs (NFkBIZ, PPP4R3B, EGR3 and IL-1β) were selected for qRT-PCR analyses. The results of the qRT-PCR showed similar trends to those observed in the microarray data. In detail, the expression levels of TCONS_00004393, NR_029386, NFkBIZ, EGR3 and IL1β were significantly increased in AG group than those in the IG and Ctr group (P<0.05,respectively; Fig 2C,D). The expression level of ENST00000566457 was significantly increased both in AG and IG group than that in the Ctr group(P<0.05,respectively). However, the expression levels of NR_026756, TCONS_00003286 and PPP4R3B were significantly decreased both in the AG and IG group than those in the Ctr group (P<0.05,respectively) (Fig 2).

### GO Analysis

GO analysis was performed to gain insight into the potential functions of gout patients in the PBMCs. Differentially expressed mRNAs from the micoarray analysis were classified into different functional categories based on the bilogical processes (BP) of the gene ontology. The number of significantly enriched GO terms that indicated upregulated mRNAs in the AG and IG groups were 946 and 1068, respectively. However, only 72 of these GO terms were shared between the AG and the IG group. In contrast, 245 and 397 GO terms of down-regulated mRNAs were enriched in the AG group and the IG group, respectively, but only 31 of these GO terms were enriched in both group. The GO analysis showed that compared with the healthy control group, the up-regulated mRNAS in the AG group were mainly involved in “nucleosome assembly”, “chromatin silencing”, “nucleosome organization”, “chromatin assembly”, “chromatin assembly or disassembly”, and “negative regulation of gene expression, epigenetic”; the down-regulated mRNAs were mainly involved in “RNA biosynthetic process”, “regulation of nucleic acid-templated transcription”, “nucleic acid metabolic process”, “regulation of transcription, DNA-templated”, and “regulation of RNA biosynthetic process”, etc (Fig 3A,B; the top10 GOs). Conversely, compared with the healthy control group, the up-regulated mRNAs in the IG group were significantly enriched in “nucleosome assembly”, “chromatin assembly”, “nucleosome organization”, “chromatin assembly or disassembly”, and “DNA packaging”; the down-regulated mRNAs were pimarily involved in “bicarbonate transport”, “cytokine-mediated signaling pathway”, “oxygen transport”, “chemokine-mediated signaling pathway”, and “cell surface receptor signaling pathway”, etc (Fig 3C,D; the top10 GOs).

**Fig 3.**
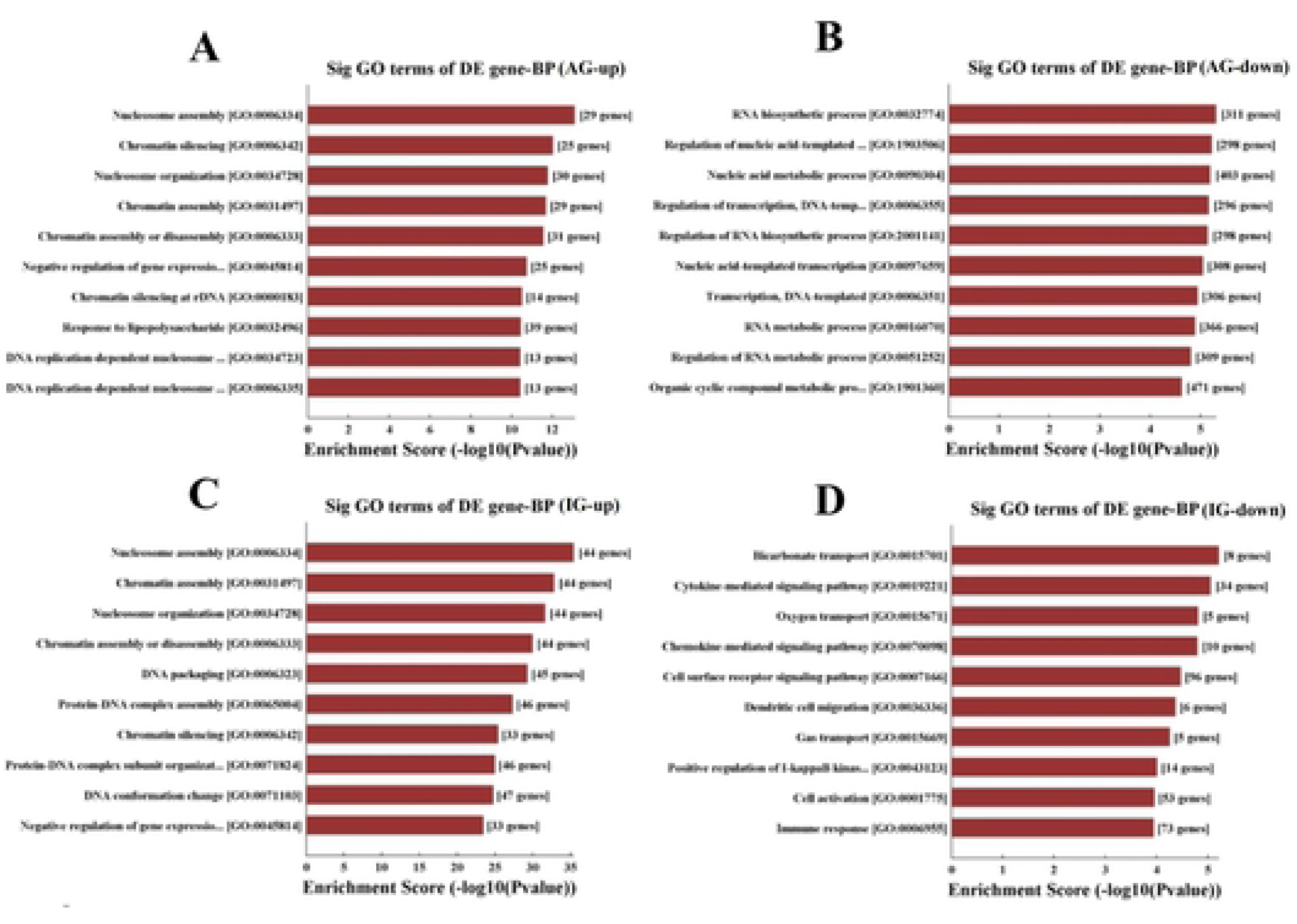
Gene ontology analyses of differentially expressed mRNAs according to biological process (BP). In GO analysis, a total of 1191 and 1465 differentially expressed mRNAs were chosen in AG and IG group, respectively. The column graphs represent the enrichment of these mRNAs, and the (− lg^P^) value has a positive correlation with GO. **A and C.** The top 10 GOs that were up-regulated in the AG and IG patients compared to the healthy controls. **B and D.** The top 10 GOs that were down-regulated in the AG and IG patients compared to the healthy controls. AG represents acute gout, IG represents intercritical gout.

### KEGG Pathway Analysis

KEGG Pathway analysis was used to investigate the involved biological pathways of the differentially expressed mRNAs. The numbers of pathway terms that the up-regulated mRNAs in the AG and IG groups were 30 and 40, respectively. Remarkably, 19 of these pathway terms were shared between the two groups. In contrast, 20 and 21 pathway terms of down-regulated mRNAs enriched in the AG group and the IG group, respectively, and 3 of these pathway terms were enriched in both group. The KEGG Pathway analysis showed that compared with the healthy control group, the up-regulated mRNAs in the AG group mainly participated in “Systemic lupus erythematosus”, “Alcoholism”, “Salmonella infection”, “Metabolism of xenobiotics by cytochrome P450”, “Viral carcinogenesis”, “toll like receptor signaling” (hsa0460), “nod like receptor signaling” (hsa0461), “IL-17 signaling pathway”(hsa04657),etc., whereas the down-regulated mRNAs were significantly involved in “Th17 cell differentiation”, “Mannose type O-glycan biosynthesis”, “Natural killer cell mediated cytotoxicity”, “DNA replication”, “Primary immunodeficiency”, and “Purine metabolism”, etc (Fig 4A,B; the top10 pathways). However, compared with the healthy control group, the up-regulated mRNAs in the IG group were significantly associated with “Alcoholism”, “Systemic lupus erythematosus”, “Viral carcinogenesis”, “Transcriptional misregulation in cancer”, “Necroptosis”, and “Legionellosis”, whereas the down-regulated mRNAs were mainly enriched in “Cytokine-cytokine receptor interaction”, “Chemokine signaling pathway”, “Influenza A”, “Peroxisome”, and “African trypanosomiasis”, etc (Fig 4C,D; the top10 pathways).

**Fig 4.**
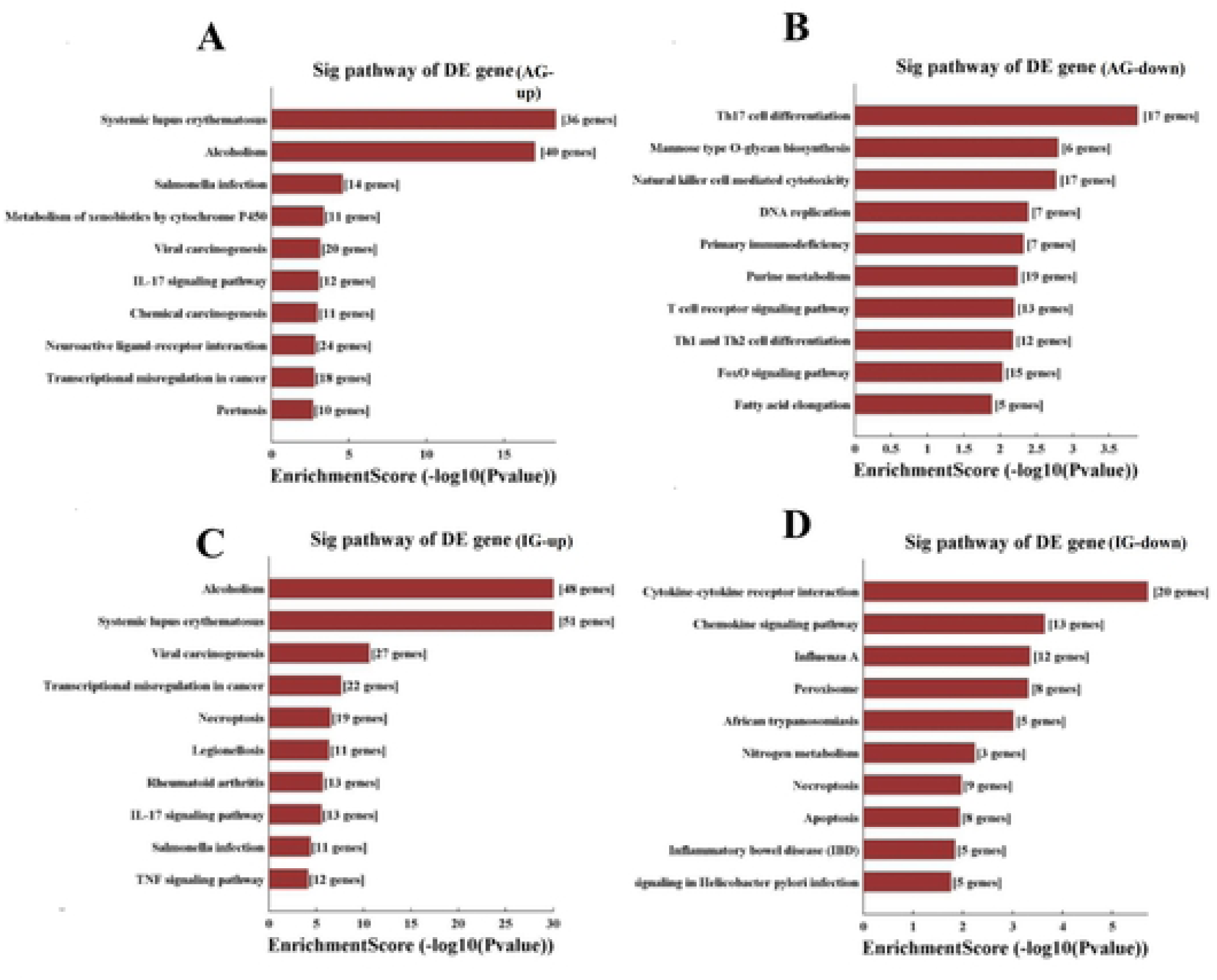
KEGG pathways analysis of differentially expressed mRNAs. In pathway analyses, a total of 50 and 61 differentially expressed mRNAs were chosen in AG and IG group, respectively. Column graphs represent the enrichment of these mRNAs, and the (−lg^P^) value has a positive correlation with pathway analyses. **A and C.** The top 10 GOs that were up-regulated in the AG and IG patients compared to the healthy controls. **B and D.** The top 10 GOs that were down-regulated in the AG and IG patients compared to the healthy controls. AG represents acute gout, IG represents intercritical gout.

### LncRNA-mRNA Co-Expression Network

The lncRNA-mRNA coexpression network may provide potential interplay between mRNAs and lncRNAs, and therefore, the differentially expressed lncRNAs and mRNAs between the AG and healthy control groups were used to draw co-expression network using the Cytoscape program. Previous studies have demonstrated that toll like receptor (TLR) signaling and NOD like receptor signaling are involved in gouty arthritis development[8]. The significantly enriched KEGG pathway analysis found that the differentially expressed lncRNAs-co-expressed mRNAs included the TLR signaling (hsa0460) and NOD like receptor signaling (hsa0461), we focus on how the lncRNAs regulate genes in the TLR signaling and NOD like receptor signaling pathways (Fig 5). The network indicated that 180 lncRNAs interacting with 7 mRNAs participated in the meaningful “NOD like receptor” signaling pathway, and 90 lncRNAs interacting with 9 mRNAs participated in the meaningful TLR signaling pathway.

**Fig 5.**
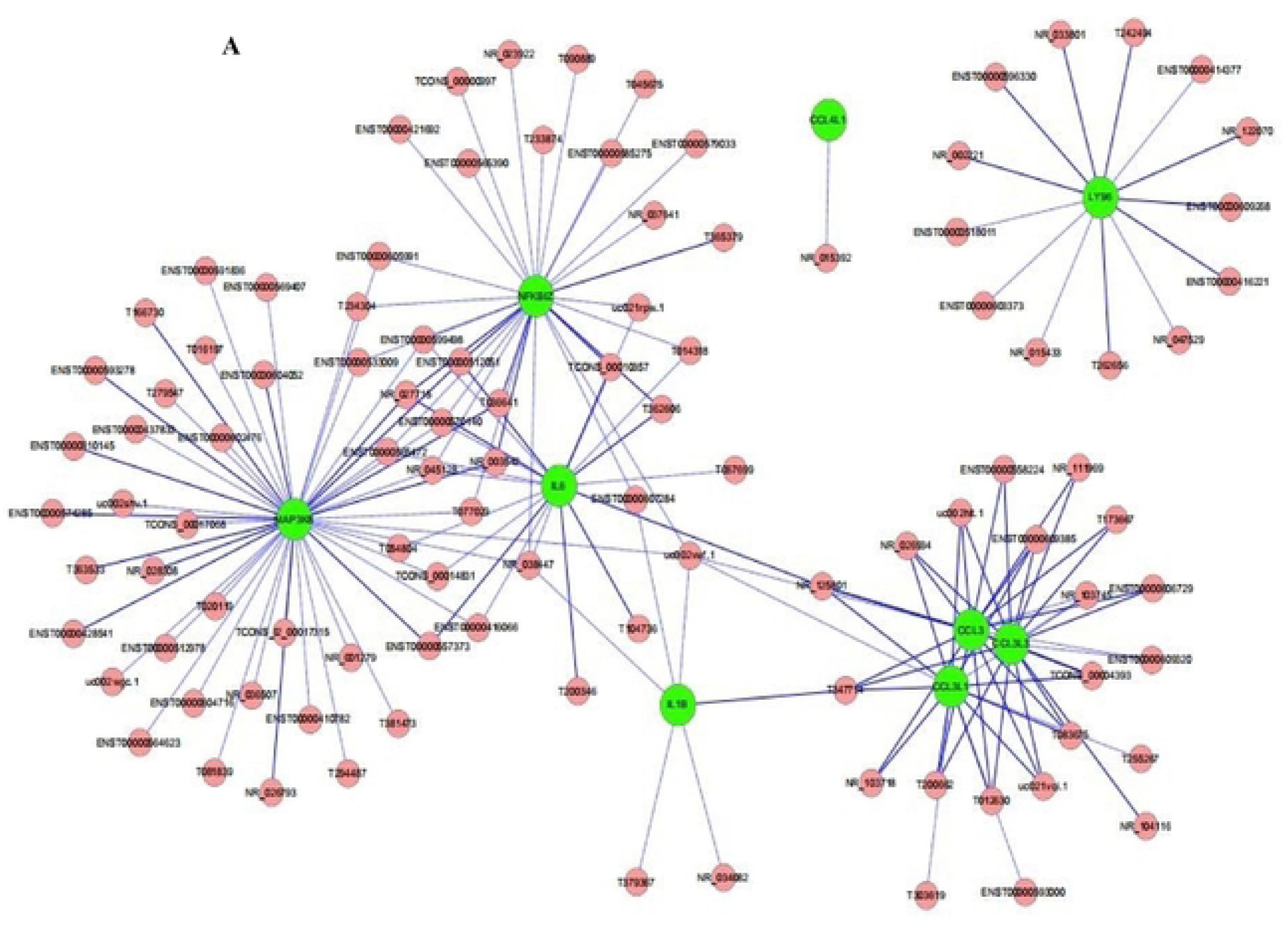
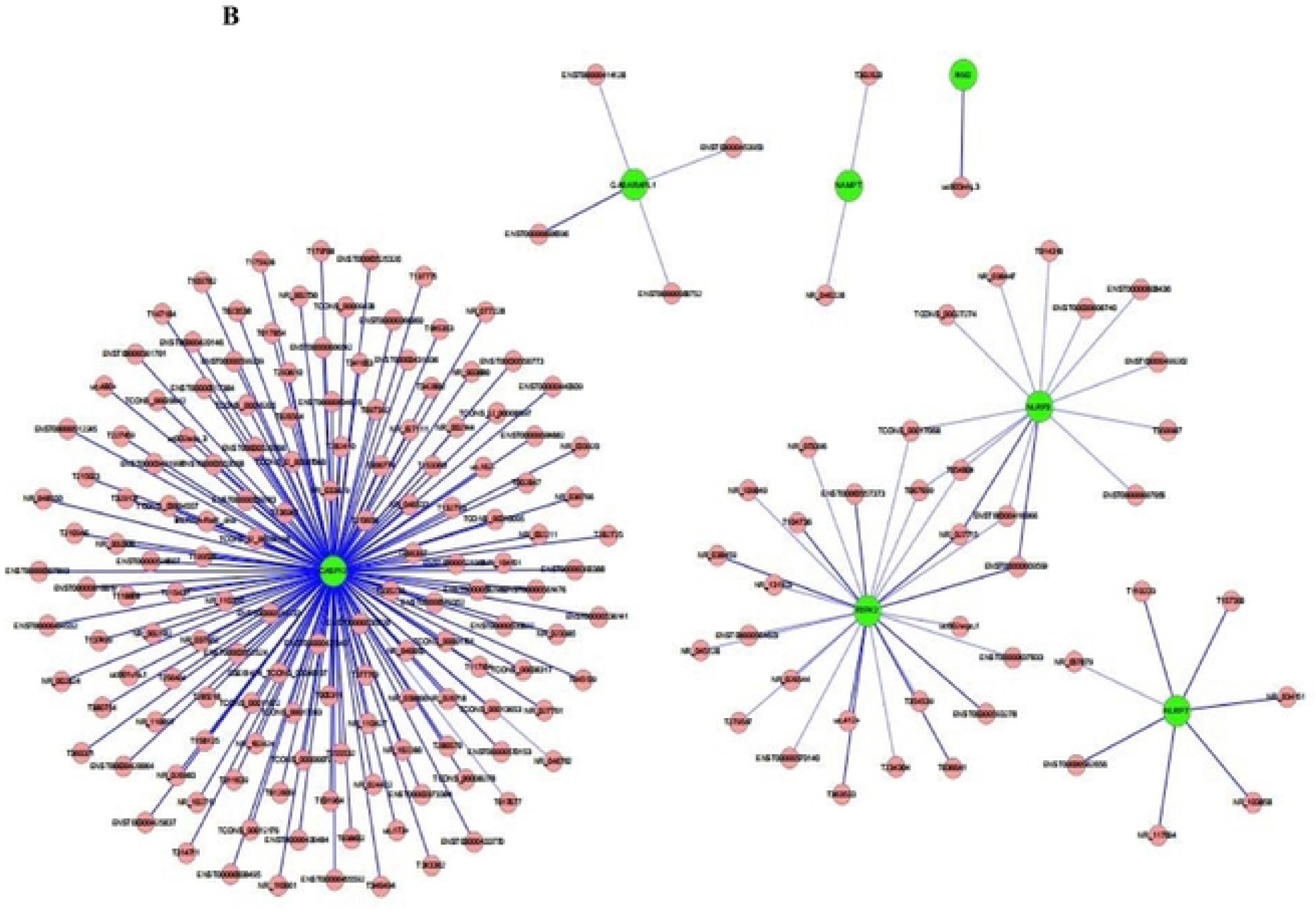
IncRNA-mRNA co-expression network. **A.** Ninety lncRNAs interacted with nine mRNAs in the meaningful “toll like receptor” signaling pathway. **B.** One hundred and eighty lncRNAs interacted with seven mRNAs in the “NOD like receptor” signaling pathway.

### Diagnostic value of the three lncRNAs for acute gout

There is an urgently needed for the identification of novel and more efficient diagnostic methods to aid in acute gout. In recent years, many studies have demonstrated that IncRNAs can serve as novel diagnostic markers for a variety of disease[9–11]. Thus, we sought to determine whether the differentially expressed lncRNAs identified in our microarray analysis could potentially serve as a diagnostic biomarkers for acute gout.

The expression of three lncRNAs, TCONS_00004393, NR_029386 and ENST00000566457, which were selected based on their significant upregulation in acute and intercritical gout patients compared to the healthy control subjects, were detected using qRT-PCR in PBMCs from 32 acute gout patients,32 intercritical gout patients and 32 healthy control subjects.The potential diagnostic value of three lncRNAs for acute gout was evaluated using healthy subjects and intercritical gout patients as control subjects, respectively. The potential diagnostic value of the three lncRNAs for intercritical gout was evaluated using healthy subjects as control subjects.

The data showed that TCONS_00004393, NR_029386 and ENST00000566457 were all significantly elevated in the acute gout patients compared to the intercritical gout and the healthy control subjects (P<0.01,respectively; Fig 6A,B,C). Expression of ENST00000566457 was significantly increased in the intercritical gout than that in the healthy control subjects(P<0.01, Fig 6A). Furthermore, receiver operating characteristic (ROC) analysis was performed to evaluate the predictive power of TCONS_00004393, NR_029386 and ENST00000566457. The area under the ROC curve (AUC) of the three candidate lncRNAs are 0.955, 0.749 and 0.961, respectively (Fig 6D) in acute gout patients compared to the healthy subjects. The AUC of the three lncRNAs are 0.870, 0.711 and 0.839, respectively (Fig 6E) in acute gout patients compared to the interciritcal gout subjects. However, in intercritical gout patients, the AUC of the three lncRNAs are 0.576, 0.469 and 0.756, respectively (Fig 6F). The data indicated that TCONS_00004393 and ENST00000566457 have high diagnostic value for acute gout.

**Fig 6.**
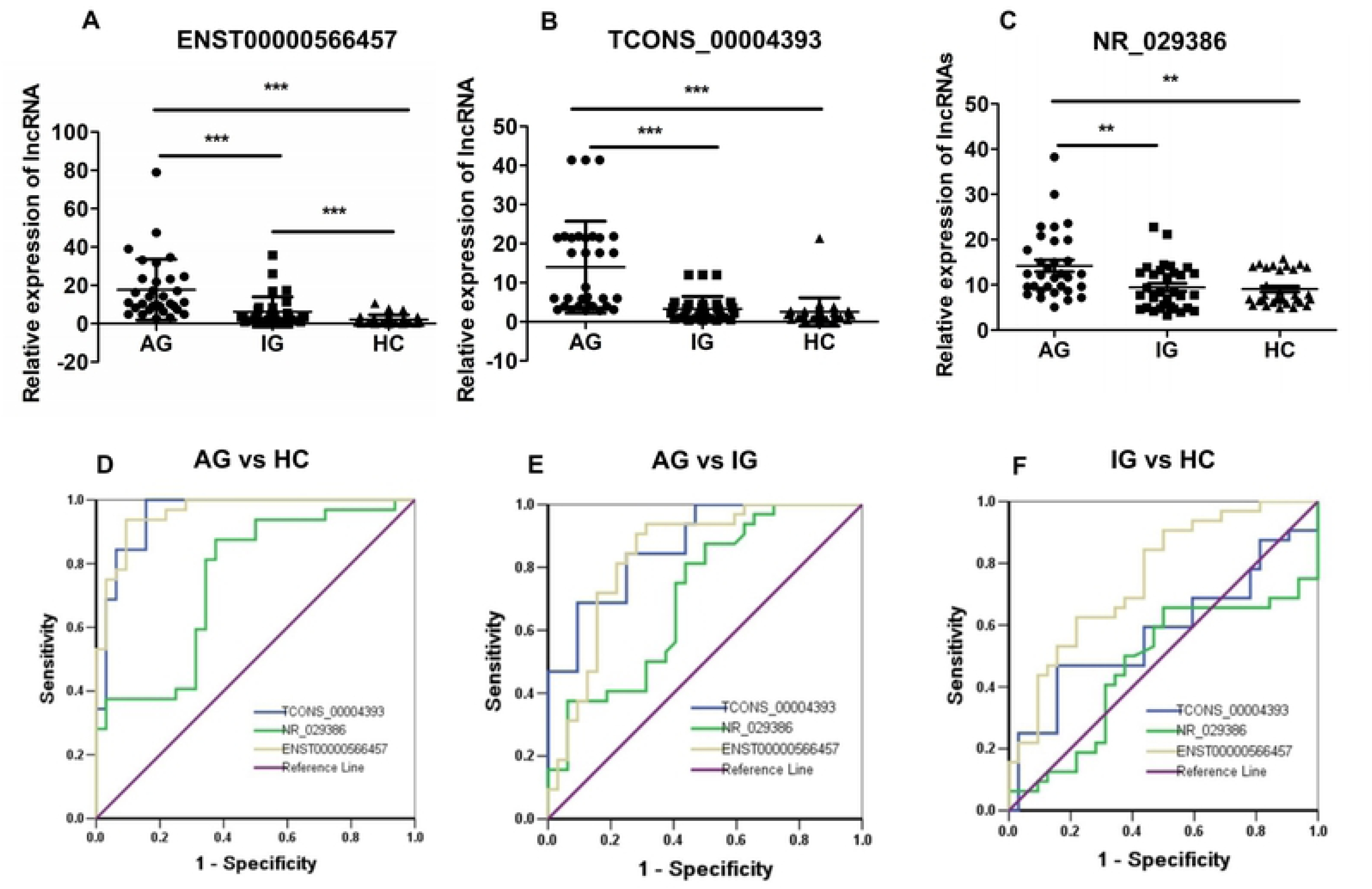
Assessment of the diagnostic value of the three lncRNAs for acute gout. **A-C.** The qPCR results of ENST00000566457 (A), TCONS_00004393 (B) and NR_029386 (C) among AG group, IG group and HC group. **D-F.** Receiver operating characteristic (ROC) curve for the three lncRNAs in AG compared to the HC (D) and IG (E), in IG compared to the HC (F). In the AG compared to the HC, the area under the ROC curve (AUC) of the three candidate lncRNAs are 0.955, 0.749 and 0.961 respectively (D); in the AG compared to the IG, The AUC are 0.870, 0.711 and 0.839 respectively (E); while in the IG compared to the HC, the AUC are 0.576, 0.469 and 0.756 respectively (F). ***P<0.001, **P<0.01. AG represents acute gout, IG represents intercritical gout, HC represents healthy control subjects.

## Discussion

Although the study of lncRNAs is a hot topic, the role of lncRNAs in the pathogenesis of gout is just beginning to be investigated. Fully exploring the role of these lncRNAs in gout would provide new insights into gouty pathogenesis. With the forefront technology of microarray analysis, we demonstrated here for the first time the expression profiles of human lncRNAs and mRNAs in patients with acute and interciritcal gout. Compared to healthy, matched to controls, acute gout patients expressed 3421 lncRNAs and 2240 mRNAs, intercritical gout patients expressed 1739 lncRNAs and 1027 mRNAs. Furthermore, GO and KEGG pathway analysis of the deferentially expressed mRNAs revealed some of the potential functions and pathways related to the pathogenesis of gout. Finally, two lncRNAs, TCONS_00004393 and ENST00000566457, were identified as the novel candidate diagnostic biomarkers for acute gout. Collectively, our results provide novel insight into the mechanism of gout development.

lncRNAs are a new class of noncoding RNAs lager than 200 nucleotides and have attracted much attention in recent medical studies. Previously, the involvement of lncRNAs in immune cell development has been reported, including dendritic cell differentiation, T cell activation, granulocytic differentiation, inhibition of T cell proliferation, Th1 cell differentiation, regulation interferon gamma (IFN-γ) expression, regulation of CD4+ Th2 lymphocyte migration, and CD4+ helper T lymphocyte differentiation[17–20]. Moreover, lncRNAs have been recognized as powerful regulators of numerous genes and pathways in the pathogenesis of inflammatory and autoimmune diseases, including SLE, RA, T1DM, MS, autoimmnue thyroid disease, Sjögren’s syndromepsoriasis, and Crohn’s disease[17,21–25].

Gout is an autoinflammatory arthropathy. Accumulating evidence indicates that genetic factors, environmental triggers and immune dysregulation might be involved in gout development, the concrete pathogenesis of gout is still unclear. The present study was undertaken to investigate the differentially expressed lncRNAs and mRNAs in gout patients compared to healthy controls. Designed for the global profiling of human lncRNAs and protein-coding transcripts, the lncRNA microarray V4.0 system was used here to screen the aberrant lncRNAs in acute and intercritical gout patients to distinguish them from those in healthy controls. Upon comparing lncRNA and mRNA expression profiles of gout patients and controls, we found that 3421 lncRNAs (2147 up-regulated and 1274 down-regulated lncRNAs) and 2240 mRNAs (1404 up-regulated and 836 downregulated mRNAs) in acute gout, 1739 lncRNAs (1046 up-regulated and 693 down-regulated) and 757 mRNAs (250 up-regulated and 507 down-regulated) in intercritical gout were differentially expressed compared to the healthy controls, and five lncRNAs and four mRNAs were further validated by qRT-PCR in 32 acute gout patients, 32 intercritical gout patients and 32 healthy subjects. Moreover, compared with the IG group, 2855 lncRNAs (1931 up-regulated and 924 down-regulated) and 1654 mRNAs (870 up-regulated and 784 down-regulated) were deregulated in the AG group (Table1). Our results improved our understanding of the molecular mechanisms of genes and lncRNAs in gout.

Go analysis and KEGG pathway analysis were performed to gain insight into the potential functions of the differentially expressed mRNAs and to improve our understanding of the mechanisms of gout development. In our results, most of the enriched up-regulated GO terms and pathway terms were shared together between the AG and the IG groups (Fig 3,4). In contrast, the significantly enriched down-regulated GO terms and pathway terms in the AG group and the IG group are quite different from each other(Fig 3,4). Previous studies have demonstrated that the TLR signaling pathway and NOD like receptor signaling pathway were both involved in the pathogenesis of gout inflammation[3,4]. Interestingly, the pathway analysis found that IL-17 signaling, Systemic lupus erythematosus, Alcoholism, Viral carcinogenesis, Salmonella infection and etc. pathways were enriched in AG and IG, except for the TLR and NOD like receptor signaling pathway. These data indicated that multiple signaling pathways associated with inflammation and metabolism were involved in the pathogenesis of gout development. These results might indicate our future research directions. It worth noting that the “Purine metabolism pathway” were depressed in the AG group, and the data indicated that acute gouty inflammation could be precision treatment by regulating purine metabolism.

Effective diagnosis of acute gout is critical for the treatment of acute gouty arthritis. To date, the discovery of MSU crystals under polarized light microscopy is the gold standard for the diagnosis of gout. However, the MSU inspection rate is very low, which make it difficult to diagnosis.Therefore, biomarkers are needed for diagnosis of acute gout, especially gout patients with no hyperuricemia. In recent years, increasing evidences have suggested that lncRNAs can serve as novel diagnostic markers for many diseases. For example, a previous report showed that a circulating lncRNA OTTHUMT00000387022 from monocytes can be used as a novel biomarker for coronary artery disease [26]. Circulating lncRNA-HULC may be a candidate serum tumour marker for the early diagnosis of gastric cancer and for monitoring its progression and prognosis [27]. However, there are no reports regarding the use of lncRNAs as biomarkers for acute gout. In this study, we found 3 lncRNAs, TCONS_00004393, NR_029386 and ENST00000566457, that in comparison with healthy subjects were significantly aberrantly expressed in PBMC samples from patients with acute gout patients. ROC analysis showed that the AUC of TCONS_00004393 and ENST00000566457 were more than 0.9, while that of NR_029386 was lower than 0.8, which indicates that TCONS_00004393 and ENST00000566457 may function as more promising candidate biomarkers for acute gout diagnosis. However, a larger sample size is needed to confirm our results. It is well known that if the AUC is over 0.9 that indicated diagnostic tests are excellent. So, TCONS_00004393 and ENST00000566457 might be effective diagnostic biomarkers for acute gout.

In summary, our study provides comprehensive lncRNA and mRNA profiles for acute gout and interciritcal gout patients. The current study provides the first demonstration that the interplay between lncRNAs and mRNA may be involved in the pathogenesis of gout, especially acute gout. More importantly, we found that two lncRNAs, TCONS_00004393 and ENST00000566457, have the potential to be novel diagnostic biomarkers for acute gout.

## Acknowledgements

This work was supported by the National Science Foundation of China (81974250); Post-Doctor Research Project, West China Hospital, Sichuan University (2018HXBH017).

## Author Contributions

Yu-Feng Qing. and Quan-Bo Zhang designed the project and wrote the paper. Jian-Xiong Zheng, Yi-Ping Tang, Fei Dai and Zeng-Rong Dong conducted the experiments. All authors reviewed and approved the final manuscript.

